# Phage-Antibiotic Combination: An Effective Method for Eradication of *Staphylococcus aureus*

**DOI:** 10.1101/2023.03.27.534482

**Authors:** Archana Loganathan, Prasanth Manohar, Ramesh Nachimuthu

## Abstract

Increasing antibiotic resistance poses a serious threat, especially in patients admitted to ICUs. The use of phages in combination with antibiotics as compassionate therapy has become a choice of treatment for pan-drug-resistant bacteria. Here, we studied the cumulative effect of phages with four antibiotics, fosfomycin, ciprofloxacin, vancomycin and oxacillin using three different treatment orders against *S. aureus*. The antibiotic disc synergy method showed that the plaque size of the phage increased in the subinhibitory antibiotic zone. The sub-inhibitory antibiotic amended in the agar media showed that the plaque size increased between 0.25 μg/mL and 1 μg/mL of antibiotics. It increases from 0.5 ± 0.1 mm (phage-alone control) to 4 ± 0.2 mm, 1.6 ± 0.1 mm, and 1.6 ± 0.4 mm with fosfomycin, ciprofloxacin, and oxacillin, respectively. Checkerboard analysis showed that phages and antibiotics were synergistic with the FIC index of less than 0.5. So, phage-antibiotic combination treatment appeared to be effective. However, the highest efficiency was observed when the antibiotics were administered after phage treatment. A maximum of 39.4-, 39.4-, and 37.0-fold reduction relative to untreated bacterial culture was observed with fosfomycin, oxacillin, and ciprofloxacin. Vancomycin antibiotic had a least 14.7-fold reduction. Finally, our findings emphasize the potential benefits of phage-antibiotic combination therapy compared with phage-alone therapy to treat *S. aureus* infections.

## 1. Introduction

Antibiotic resistance in bacteria occurs because of modifications or adaptations to the drugs used for treatment, and in a few bacteria, it can also occur because of the intrinsic mechanisms [1]. A decline in the efficacy of antibiotics for the treatment of drug-resistant bacteria has re-established phage therapy as one of the recent clinical practices. Phage therapy involves the use of bacterial viruses (bacteriophages) to treat multidrug-resistant bacterial infections [2]. An antimicrobial review conducted in 2016 estimated that the death due to antimicrobial resistance (AMR) would increase to 10 million each year by 2050 [3]. *Staphylococcus aureus* is a notorious pathogen that tends to rapidly develop resistance. The evolution of antibiotic resistance in *S. aureus* has been well documented [4] and it is known to be one of the top pathogens causing more than 100,000 deaths in 2019 [1]. These examples suggest that *S. aureus* has grown out to be a deadly pathogen that requires immediate attention and an alternative treatment strategy.

Phage therapy has recently become a more pronounced treatment option in many developed countries [5]. Most clinical reports on phage treatment have shown promising therapeutic outcomes [6]. Additionally, clinical studies have demonstrated encouraging results in treating osteomyelitis [7], bone and joint infections [8] [9], and pulmonary infections [6]. Most clinical studies have reported that the combination of phages and antibiotics is more effective than phage treatment alone [10,11]. Phage and antibiotic combination treatment describes the use of phage and sub-inhibitory concentrations of antibiotics that can work in synergy, whereby the efficacy of the sum is greater than the individual effect. The combination of phages and antibiotics has been proven to be clinically significant owing to some therapeutic benefits that outnumber phage-alone treatment. For example, a sub-inhibitory concentration of antibiotics and phages in combination was found to increase the burst size of phages [12], eradicate biofilms [13], and enable an evolutionary trade-off [14] [15] [16]. In clinics, this type of therapy is chosen after confirming the synergistic effect of the antibiotic and phage, because a few antibiotics that operate synergistically with one phage do not have the same effect on others, resulting in an antagonistic combination. For instance, rifampicin, in combination with staphylococcal phage sb-1, is antagonistic [17], whereas it shows synergism with phage SAP-26 [18]. Phage therapy against *S. aureus* is well documented [19-22]. Commonly reported antibiotics used in combination with staphylococcal phages to include penicillin [23], gentamycin [24], rifampicin [18], linezolid [25], ciprofloxacin [26] and vancomycin [27]. There is numerous research pertaining to phage-antibiotic synergy (PAS) in the *S. aureus* [28]; however, studies on the treatment order in PAS are limited. The PAS approach reported in this study should be able to address the eradication efficiency of phage-antibiotic combination treatment.

We previously reported that phage vB_Sau_S90 was effective against 206 clinical isolates of *S. aureus* [29]. The current (preliminary) study aimed to investigate the efficiency of the phage in combination with four different antibiotics (fosfomycin, ciprofloxacin, oxacillin, and vancomycin) and to identify the most effective treatment strategy. Furthermore, the change in plaque size was analyzed by amending the DAOL (Double agar overlay) plate with a range of sub-inhibitory concentrations of antibiotics. We also studied the synergism between different combinations of phage-antibiotic from lower to higher concentrations using checkerboard analysis. Finally, the sub-inhibitory concentration of the antibiotic and the phage was determined using three different treatment orders (phage treatment followed by antibiotic treatment (PRE), simultaneous phage and antibiotic treatment (SIM), and antibiotic treatment followed by phage treatment (POS)).

## 2. Materials and Methods

### 2.1 Bacteria, bacteriophage, and culture conditions

A clinical *S. aureus* strain, SA-90, and phage vB_Sau_S90 used in this study were previously reported [29]. Bacterial isolate SA-90 was originally obtained from a diagnostic laboratory in Chennai, India [24]. The isolate was obtained from a pus sample, and the antibiotic resistance analysis by PCR showed that the isolate harboured the *mec*A gene, thus classified as methicillin-resistant *S. aureus* (MRSA). Bacterial cultures were maintained in Brain Heart Infusion broth (HiMedia, India) supplemented with agar when required. The bacterial culture was maintained at 37 □. Phage vB_Sau_S90 was isolated from a hospital sewage sample [29]. Throughout the study, the DAOL was performed using soft agar of 0.45% (composition: LB broth: 25 g/L, bacteriological agar-agar: 0.45%). Transmission Electron Microscopy analysis revealed that the phage exhibited a *Podoviridae* morphology (Supp. Fig 1). Characteristic features of the phage, including its life cycle and time-kill analysis, have been reported previously [29].

### 2.2 Checkerboard analysis

The outcome of the phages and antibiotics combination was analyzed using the checkerboard method. Briefly, the effect was assessed using phage vB_Sau_S90 and antibiotics fosfomycin, ciprofloxacin, oxacillin, and vancomycin. Phage dilutions from 10^2^ to 10^9^ PFU/mL were diluted along the abscissa and marked as plate A, and antibiotics from 64 to 0.125 μg/mL were diluted along the ordinates and marked as plate B. The phage dilutions from plate A were transferred to plate B in their respective wells in 96-well titer plates (HiMedia, India). The inoculum of the test bacterial isolate SA-90 was prepared by diluting the overnight culture to 0.5 McFarland turbidity, and 5 μl of the inoculum was added to each well of a 96-well titer plate that contained the mixture of phage and antibiotic. The effect of the combination was determined by measuring the fractional inhibitory concentration (FIC) index. The FIC index was calculated as described elsewhere [30]. The effect was interpreted based on the FIC index, which can be interpreted as synergistic (FIC < 0.5), additive (2 > FIC ≤ 0.5), or antagonistic (FIC ≥ 2).

The percentage of bacterial reduction at different combinations was calculated by measuring the optical density (OD = 600 nm) and converting the results to a percentage reduction value ((OD of treated/OD of untreated) × 100%). Antibiotic-only (128-0.25 μg/mL) and phage-only (10^2^ to 10^9^ PFU/mL) and bacteria-only were maintained as a control. The antibiotics that showed synergism were considered for further analysis.

### 2.3 Disk Diffusion phage synergy test

The synergy test was performed as previously described [31]. Briefly, a double agar overlay (DAOL) was performed using phage vB_Sau_S90 (1x10^2^ PFU/mL) and an antibiotics disk was placed in the centre. Antibiotic concentration was based on the standard antibiotic disk concentration outlined in the Clinical and Laboratory Standards Institute (CLSI) guidelines. The phage concentration was selected such that neither too many nor too few plaques were produced on the double agar overlay (DAOL) plate without an antibiotics disk and the same concentration of phage was used for the analysis of plaque size with the antibiotic disk. The phage concentration may vary for each phage, depending on the size of the plaques, and may need to be adjusted accordingly. The following antibiotics that showed synergistic activity in checkerboard analysis were tested: fosfomycin (200 μg), ciprofloxacin (5 μg), oxacillin (30 μg), and vancomycin (30 μg) (HiMedia, India). The plates were incubated at 37 °C for 16 hrs. Changes in plaque size were observed near the sub-inhibitory concentration zone. Note that our definition for sub-inhibitory concentration in the disc diffusion DAOL test is the point immediately near the periphery of the zone of inhibition where bacteria is not inhibited or killed but provides a measure of plaque size. DAOL plate with an empty Whatman filter paper disk (no antibiotics) was maintained as a control.

### 2.4 Agar plate phage synergy test

The agar plate phage synergy test was conducted to assess the change in plaque size at various sub-inhibitory concentrations of antibiotics. Minimum inhibitory concentration was found to be 8 μg/mL, 16 μg/mL, 8 μg/mL, and 2 μg/mL for fosfomycin, ciprofloxacin, oxacillin, and vancomycin, respectively. The antibiotic concentrations below the MIC (8, 4, 2, 1, 0.5, 0.25 μg/mL) were initially screened for the agar plate synergy test. The antibiotic concentration that prevented bacterial lawn formation was considered as activity by antibiotic alone (data not shown); thus, those concentrations were eliminated, and only those concentrations that did not inhibit the bacterial lawn formation and enhanced the plaque size were considered synergistic concentrations. The antibiotic concentrations, 0.25 μg/mL, 0.5 μg/mL, and 1 μg/mL were found to be synergistic for all four antibiotics (fosfomycin, ciprofloxacin, oxacillin, and vancomycin). Briefly, an agar plate synergy test was performed by adding a sub-inhibitory concentration of antibiotic to the hard agar and DAOL was performed on the antibiotic media using phage vB_Sau_S90 (1x10^2^ PFU/mL). The plates were incubated at 37 □ for 16 – 18 hrs. The effect of antibiotics on the plaque size was observed after 18 hrs. Double agar overlay without antibiotics was maintained as a control. The difference in the plaque size in each antibiotic concentration was measured manually and compared to the control. Experiments were performed in triplicates and statistical analyses were performed using the mean (SD).

### 2.5 Phage-antibiotic synergy

Time-kill analysis was conducted to assess the decrease in bacterial turbidity resulting from the combination of phage and antibiotics. Here, the combinational effect of phage-antibiotic was evaluated using three treatment orders: phage treatment followed by antibiotic treatment (PRE), simultaneous phage and antibiotic treatment (SIM), and antibiotic treatment followed by phage treatment (POS).

Briefly, to the bacterial inoculum of 0.2 at OD600 nm (bacterial burden -1 × 10^8^ CFU/mL), the phage at 10^7^ PFU/mL and the antibiotics at a concentration of 1 μg/mL were added in the PRE, SIM, POS treatment orders, independently. The bacterial turbidity was measured every 2 hrs for 22 hrs at OD600 nm. The time interval between the phage and antibiotic addition in PRE and POS treatment was 60 min, and 0 min for SIM treatment. The effective treatment order was determined by calculating the fold-reduction in bacterial growth, as described previously [32]. The effectiveness of phage-antibiotic synergy was interpreted using SIM treatment as most of the studies reported so far tested PAS by adding phages and antibiotics at the same time. The effective treatment order was interpreted by comparing all three treatment orders. The efficacy of the method was validated by comparing the percentage reduction of treatments with controls, which include phage-only (Bacteria+10^7^ PFU/mL), antibiotic-only (Bacteria+1 μg/mL), and bacteria-only. Percentage reduction was calculated as follows: (OD of untreated -OD of treatment/OD of untreated) × 100. The OD of the untreated sample represented the bacteria-only control, and the OD of the treatment represented the phage-only, antibiotic-only, and phage-antibiotic combination. Percentage reduction was calculated for all three treatments compared to the untreated bacteria-only control.

### 2.6 Statistical analysis

Data are representative of mean ± SD (n=3). The outcomes of the treatment orders were compared with the antibiotics using one-way ANOVA. The statistical difference between the treatment order of an antibiotic and the phage-alone treatment was compared using the Student’s t-test, and the treatment orders among the antibiotics were compared using a one-sample t-test. Statistical differences were evaluated at p < 0.05.

## 3. Results

### 3.1 Checkerboard analysis showed synergisms to all four antibiotics

The results of the checkerboard analysis showed that the phage and all four antibiotics in combination had a synergistic effect (FIC index < 0.5). The FIC indices of the antibiotics in combination with phage vB_Sau_S90 were 0.126, 0.072, 0.031, and 0.125 for fosfomycin, ciprofloxacin, oxacillin, and vancomycin, respectively. The combination of phages and antibiotics reduced the MIC by multiple folds compared to the individual effect. For fosfomycin, the concentration of 8 μg/mL in the individual treatments was reduced to 1 μg/mL in combination with the phage. Similarly, for the ciprofloxacin antibiotic, it reduced from 16 to 1 μg/mL, for oxacillin from 8 μg/mL to 0.25 μg/mL and for vancomycin antibiotic from 2 μg/mL to 0.25 μg/mL. The MIC of the phage alone showed a reduction at 10^8^ PFU/mL and in combinations showed a reduction at 10^5^, 10^6^, 10^4^, 10^3^ PFU/mL with fosfomycin, ciprofloxacin, oxacillin, and vancomycin, respectively. The combinatorial effect of phage and antibiotic at different concentrations in checkerboard analysis is represented as a heatmap in Fig 1.

**Figure 1.**
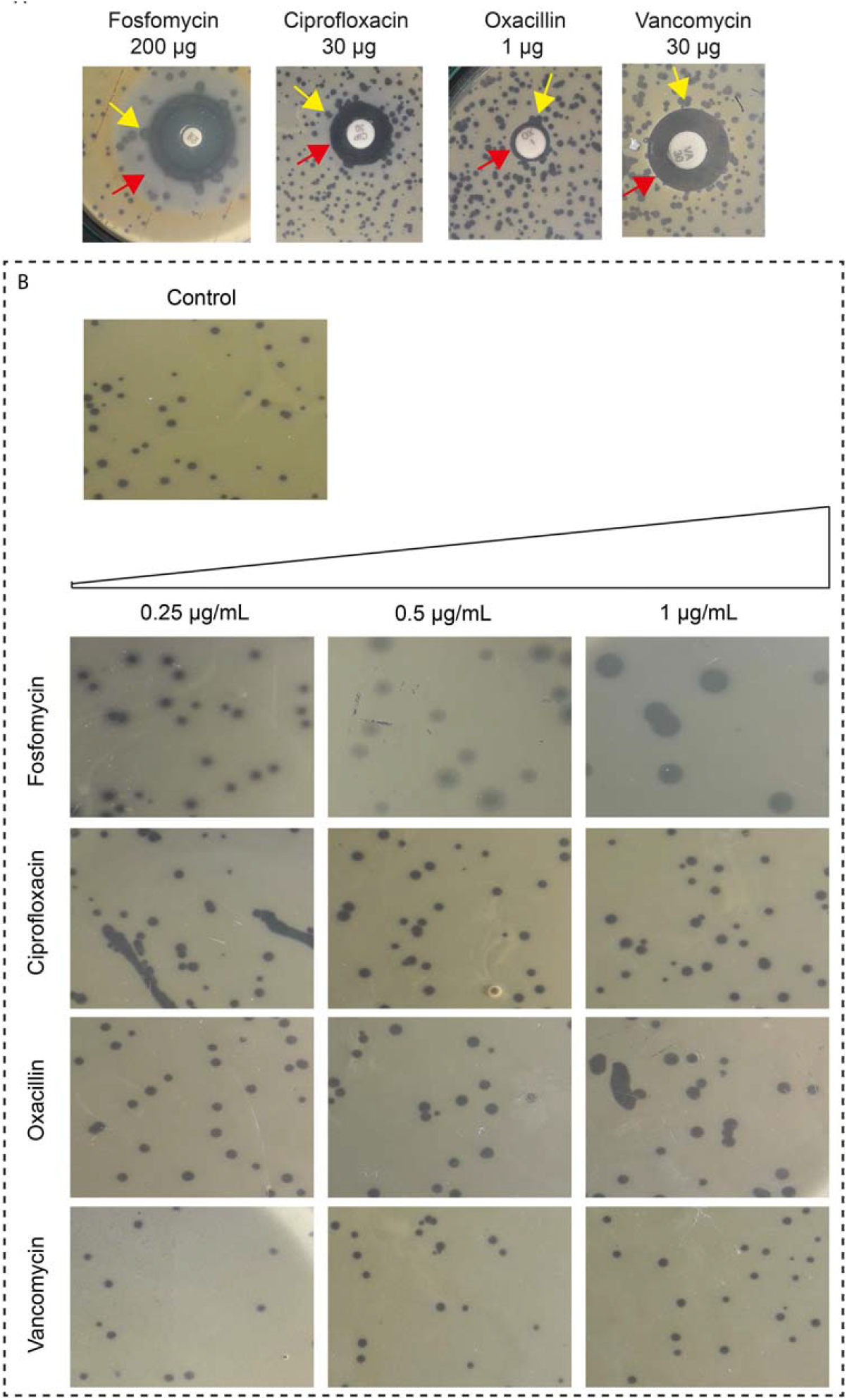
The effect of antibiotics on plaque size with fosfomycin, ciprofloxacin, oxacillin, and vancomycin: (a) *In vitro* disk diffusion synergy. The red arrow represents the subinhibitory concentration zone and the yellow arrow represents the plaque that showed variation at the zone of subinhibitory antibiotic concentration; (b) Agar plate synergy at 0.25 to 1 μg/mL.

### 3.2 Phage-antibiotic combination is better than phages and antibiotics alone

The effect of the combination (SIM) and individual (phage-alone) treatment were compared using time-kill analysis. Here, simultaneous treatment (SIM) outcome was used to determine the effect of PAS treatment, as most of the *in vitro* studies in the literature determine the effect of PAS treatment without any treatment order. Phage-antibiotic combination treatment resulted in more than an 80% reduction in bacterial growth, whereas phage-alone showed a 76.6% reduction. In antibiotic-treated cells, the reduction was 58.5%, 64.5%, 64.8%, and 68.2% for fosfomycin, ciprofloxacin, oxacillin, and vancomycin, respectively. Under combinatorial treatment, fosfomycin efficiency increased 2.3 × times compared to antibiotic treatment alone and 1.3 × times compared to phage treatment alone. The ciprofloxacin-phage combination efficiency increased 2.1 × times compared to antibiotic-alone and 1.38 × times greater than phage-alone treatment. The oxacillin-phage combination efficiency increased 2.1 × times compared to the antibiotic-alone and 1.45 × times greater than that of the phage-alone treatment. The vancomycin-phage combination efficiency increased to 1.8 × compared to that of antibiotic-alone and 1.36 × greater than that of phage-alone treatment. The outcome of the checkerboard analysis is shown as the interactive plot in Fig 2.

**Figure 2.**
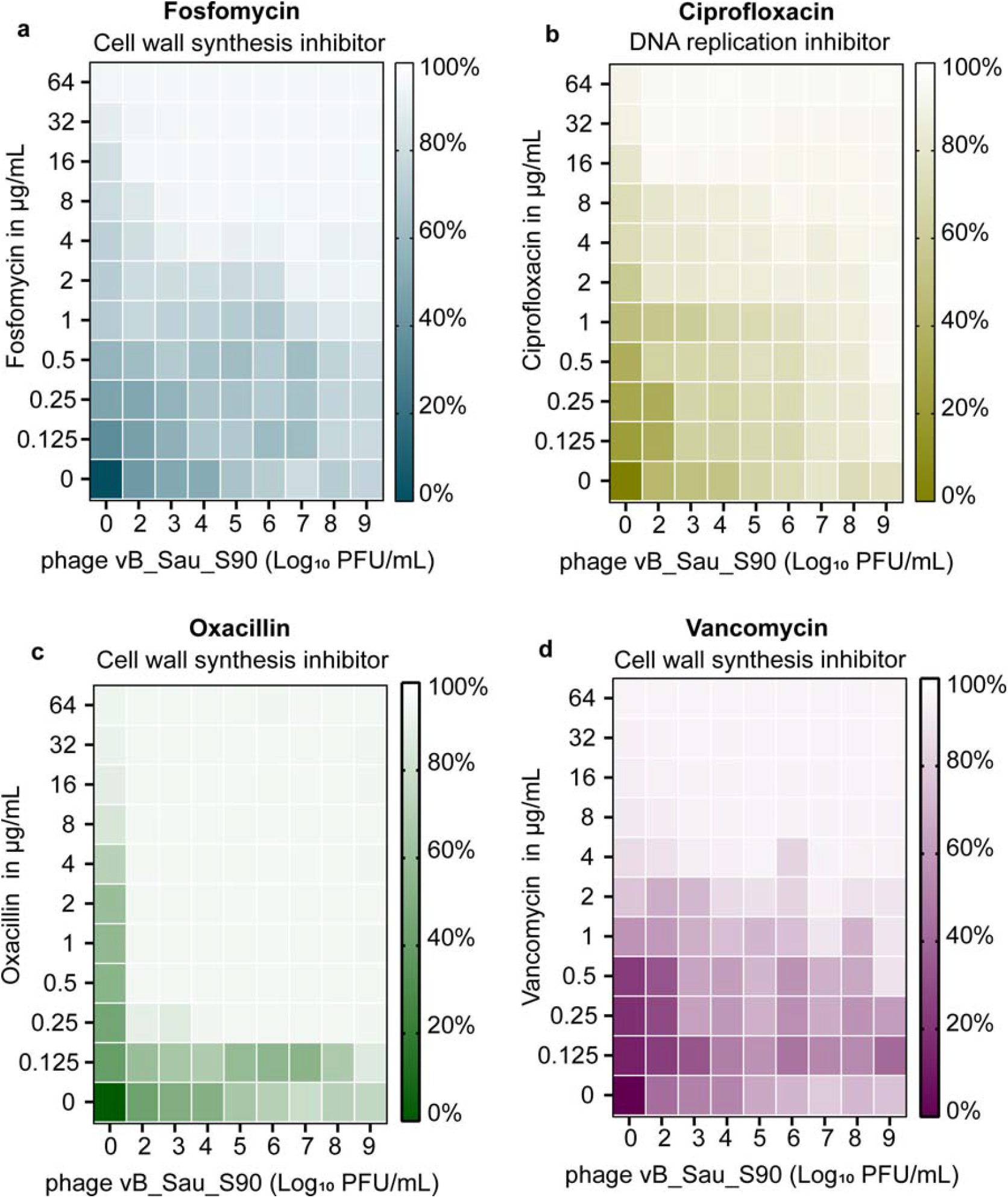
Checkerboard analysis with antibiotics along the ordinate (0.125 to 64 μg/mL) and phage vB_Sau_S90 along the abscissa (endpoint n = 16), the percentage of reduction in the bacterial turbidity is represented as a heatmap: (a) fosfomycin; (b) ciprofloxacin; (c) oxacillin; (d) vancomycin.

### 3.3 Not all antibiotics had a similar effect on the phage plaque size

We first determined the change in plaque size using various phage-antibiotic combinations. Preliminary analysis by disk diffusion method showed that plaque size was enhanced near the antibiotic zone. In particular, the effect was highly evident near the subinhibitory antibiotic zone. Change in the plaque size near the subinhibitory antibiotic zone is represented in Fig 3a. We further confirmed this effect of antibiotics on the plaque size by agar plate synergy test, which showed that the synergistic sub-inhibitory concentrations were 0.25 μg/mL, 0.5 μg/mL, and 1 μg/mL for all four antibiotics studied. The plaque size in the agar plate synergy test is shown in Fig 3b. The agar plate synergy test showed that plaque size increased gradually with increasing subinhibitory concentration of synergistic antibiotics. Fosfomycin showed the largest plaque of all four antibiotics studied, showing 9x times larger plaque compared to the phage-only treatment. A maximum plaque size of 4 ± 0.2 mm was observed at 1 μg/mL with fosfomycin, while other antibiotics had a minimal effect on the plaque size showing 3.2x times at 1 μg/mL with ciprofloxacin (1.6 ± 0.1 mm) and oxacillin (1.6 ± 0.4 mm), respectively. No significant changes were observed with vancomycin at any of the studied concentrations (Table 1).

**Table 1:** Plaque size variation with the increasing subinhibitory antibiotic concentration (0.25 to 1 μg/mL) of antibiotics. Data are presented as the mean ± SD (n = 3).

| Antibiotic concentration | 0.25 µg/mL | 0.5 µg/mL | 1 µg/mL |
| --- | --- | --- | --- |
| **Fosfomycin** | 2.6 ± 0.4 mm | 3.5 ± 0.5 mm | 4 ± 0.2 mm |
| **Ciprofloxacin** | 1.2 ± 0.2 mm | 1.3 ± 0.1 mm | 1.6 ± 0.1 mm |
| **Oxacillin** | 1.1 ± 0.2 mm | 1.6 ± 0.4 mm | 1.6 ± 0.4 mm |
| **Vancomycin** | 0.5 ± 0.1 mm | 0.5 ± 0.1 mm | 0.5 ± 0.1 mm |

**Figure 3.**
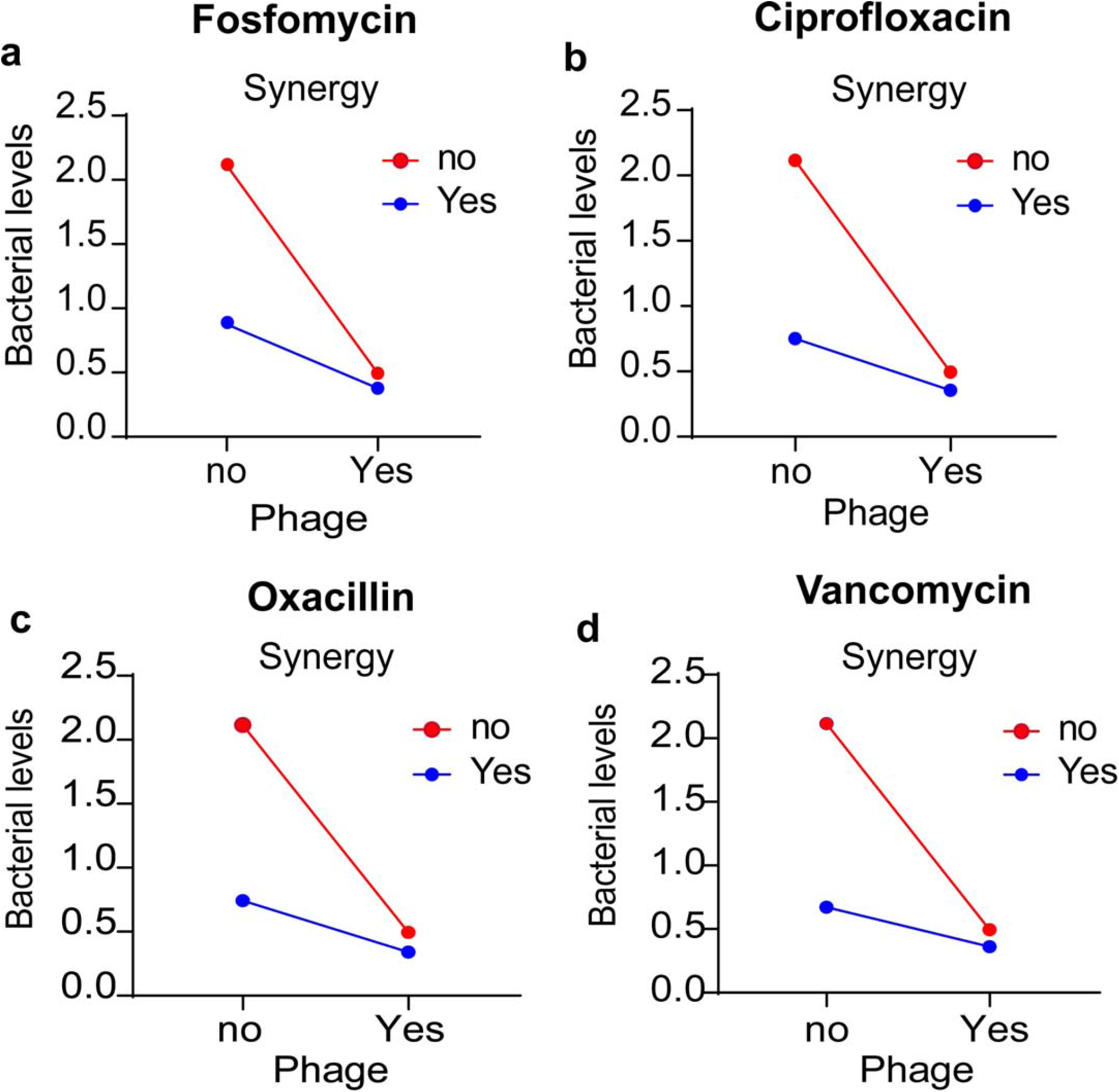
Interactive plot of the checkerboard analysis: (a) fosfomycin; (b) ciprofloxacin; (c) oxacillin; (d) vancomycin.

### 3.4 Phage followed by an antibiotic treatment was effective to other treatment orders

The efficiency of PAS treatment order analyzed by time-kill analysis showed that the order of phage administered determines the outcome of PAS treatment. The efficiency of treatment orders decreased in the order: phage followed by antibiotic treatment, antibiotic followed by phage treatment, and simultaneous treatment (POS > PRE > SIM). The fold reduction for phage followed by antibiotic treatment was 39.40-, 37.05-, 39.40-, and 14.75-fold with fosfomycin, ciprofloxacin, oxacillin, and vancomycin antibiotics, respectively. Similarly, for the antibiotic followed by phage treatment, there were 8.93-, 7.47-,9.19-, and 12.33-fold reductions with fosfomycin, ciprofloxacin, oxacillin, and vancomycin, respectively. The least reduction was seen in simultaneous treatment, ranging from 5.68-, 6.26-, 6.48-, and 6.27-fold with fosfomycin, ciprofloxacin, oxacillin, and vancomycin antibiotics, respectively. The statistical analysis showed that different treatment orders among the antibiotics were significantly different with a probability value less than 0.001 (statistical difference, p<0.05).

The fold-reduction in the bacterial growth among the antibiotics and treatment order is represented as a box plot in Fig 4.

**Figure 4.**
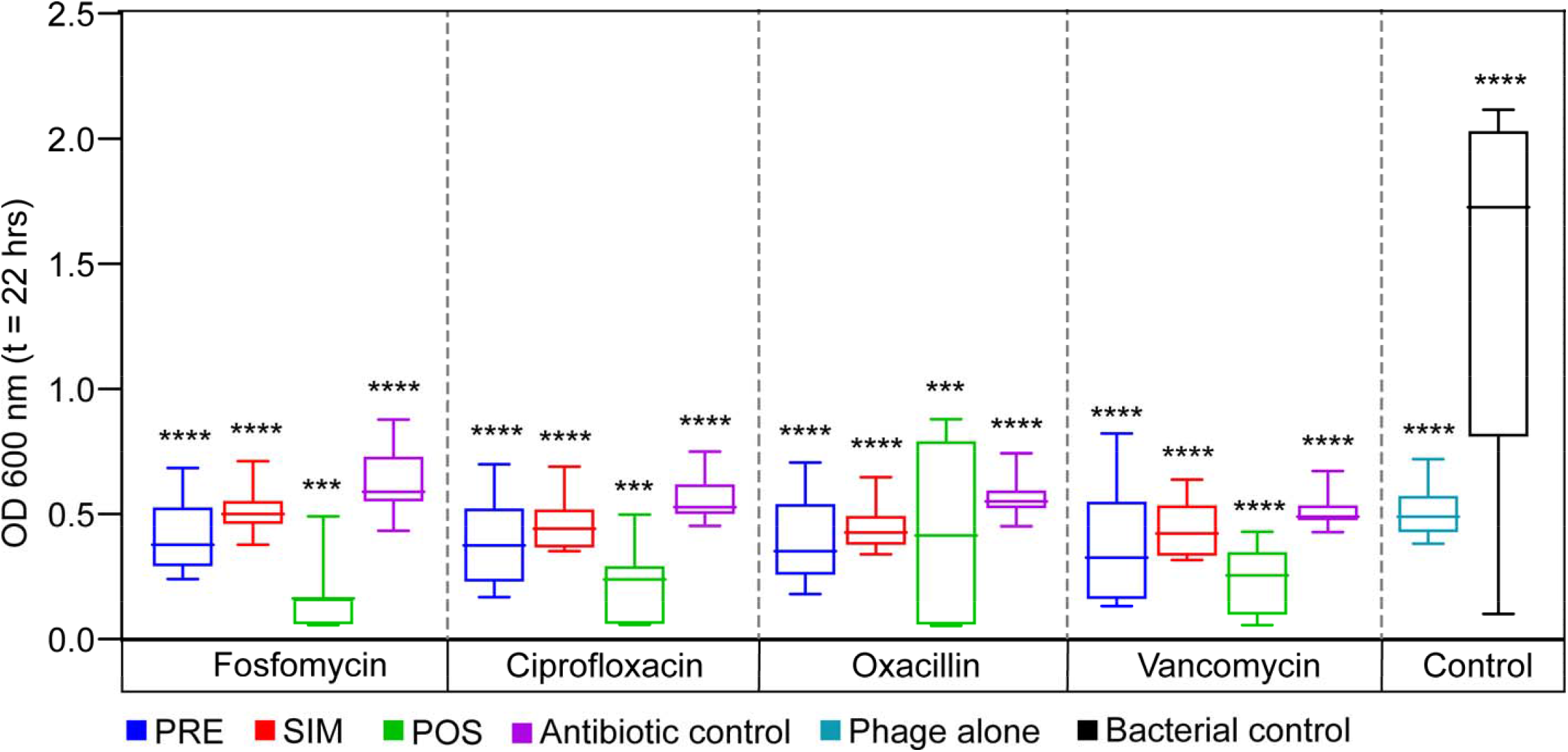
The fold reduction in the bacterial growth of three different treatment orders of four different antibiotics is presented as a box plot (endpoint at OD_600 nm_; t = 22 h). Data are presented as mean ± SD (n = 3). A one-sample t-test was used for statistical analysis (treatment order among the antibiotics); p < 0.05, (^****^p = <0.0001).

### 3.5 Effect of PAS within and among the antibiotic mechanism of action

The effect of the mechanism of action showed that the average fold reduction (average of all three treatment orders) was high for cell-wall synthesis inhibitors, fosfomycin and oxacillin, showing 18.00- and 18.36-fold, respectively. DNA synthesis inhibitors (ciprofloxacin), showed 16.93-fold bacterial reduction; however, the fold reduction was found to be higher than that of vancomycin by 5.82-fold. The effect of PAS within the cell-wall synthesis inhibitor showed that oxacillin resulted in the highest fold-reduction of bacteria.

## 4. Discussion

These results show that the capacity of phages and antibiotics in combination to eliminate the bacterial burden is superior to that of phage-only and antibiotic-only treatments. Our finding concerning the effect of the phage and antibiotic combination against *S. aureus* highlights the primary finding of PAS methods in comparison with phage-only treatment. Our study revealed that 1) plaque size variation with the addition of antibiotics is not a direct measure of synergism; however, antibiotics induced a plaque size enhancement with few antibiotics, 2) not all antibiotics with the same mechanism of action exhibited similar outcomes, and 3) the treatment order in PAS is an important determinant of treatment success, in which phage followed by antibiotic treatment was more effective than other PAS treatment orders.

The increase in plaque size with the addition of antibiotics is an important factor to be noted in this study. Our disc synergy test showed that the impact on plaque size was clearly observed near the subinhibitory zone. This is based on the principle that the antibiotic from the disc diffuses gradient into the surrounding media based on the concentration of antibiotics in the disc and creates a sub-inhibitory zone beyond the inhibitory zone. However, the exact concentration at the sub-inhibitory zone cannot be determined, thus, when the different sub-inhibitory concentrations of antibiotics were amended in the media showed that the maximum plaque size variation was observed at 1 μg/mL and remained constant thereafter for all four antibiotics studied. The plaque size in the phage and antibiotic groups showed the highest plaque size variation (increase) with the fosfomycin antibiotic at 1 μg/mL. The mechanisms of action of the antibiotics used in the study were cell wall synthesis inhibitors (fosfomycin, oxacillin, and vancomycin) and a DNA replication inhibitor (ciprofloxacin). All antibiotics used in this study showed a synergistic effect in combination with phage vB_Sau_S90; however, a varying fold reduction in bacterial growth was observed between the antibiotics with the same phage. We hypothesized that such variation within the same phage could be the outcome of the point of action of antibiotics that support phage replication. To better understand the synergism and role of antibiotics in PAS treatment, we compared the combinatorial effects, phages-alone, and antibiotics-alone based on the mechanism of action of antibiotics. In our study, the plaque size increased prominently when phages were incubated along with fosfomycin antibiotics; however, no such large variation was observed with oxacillin and vancomycin antibiotics, which also share the same mechanism of action. Variations among the same mechanistic classes can be attributed to their point of action. When examining the exact mechanism of action of each antibiotic, fosfomycin acts in the initial step of cell wall peptidoglycan synthesis, inhibiting phosphoenolpyruvate synthetase [33]. In contrast, oxacillin inhibits binding to the PBP2a protein, which is the third or last stage of cell wall synthesis, while vancomycin binds to the D-Ala terminal and prevents cross-linking [34] [35]. This shows that, within the same class of antibiotics, the point and time of action can differ, which can influence the variation in the effect of antibiotics on the plaque.

Similarly, in order to understand whether a change in plaque size is a measure of synergism between phages and antibiotics, we compared the plaque size change with antibiotics and their respective fold reduction in PAS. In both assays, we observed that the addition of fosfomycin either as a disk or when amended into the DAOL plate resulted in a drastic increase in plaque size in sub-inhibitory concentration ranges. For other antibiotics, moderate-to-no change in plaque size was observed. To determine whether increasing plaque size with antibiotics is an indirect measure of synergism, we compared the results of plaque size with the average fold reduction of bacterial growth during three treatment orders. A high fold reduction in bacterial growth was observed for the oxacillin and fosfomycin antibiotics, and the highest plaque size was observed only in the fosfomycin antibiotic disk. Although fosfomycin produced the largest plaque among the antibiotics studied, it showed the second-highest fold reduction in bacterial growth in the PAS test. Similarly, oxacillin produced a moderate change in plaque size and produced a high-fold bacterial reduction in the PAS test. In contrast, vancomycin did not have any effect on plaque size and similarly showed the least effect in combinatorial treatment. This suggests that plaque size production is independent and varies with the antibiotic; thus, it cannot determine the combinatorial effect in PAS analysis.

Preliminary and confirmatory tests were performed to determine the effect of antibiotics on plaque formation. These methods could have limitations for antibiotics with low diffusion ability. For instance, we hypothesized that the poor plaque size variation with vancomycin antibiotics could be due to their low diffusion. Vancomycin antibiotic discs tend to have low diffusion in agar media; thus, they are not preferred for disk diffusion assays, as recommended by the CLSI guidelines (Clinical Laboratory Standard Institute). However, amending antibiotics in agar had the same effect as the disk method. This indicates that the point and time of action may be determinants of plaque variation and not the mechanism of action. Similarly, when examining the effect of antibiotic susceptibility of the isolate, our study showed that the bacterial isolate SA-90 was susceptible to fosfomycin and vancomycin antibiotics. On comparing plaque size variation with the susceptible antibiotic, fosfomycin induced plaque size variation, whereas vancomycin did not have any effect, suggesting that plaque size cannot be induced by susceptible antibiotics. A study conducted by Manohar et al. showed that no plaque variation was observed in any of the antibiotic studies; however, the study showed that the reduction in bacterial burden with the same antibiotics in the time-kill analysis was significantly high [30]. In comparison with this study, it can be demonstrated that plaque size does not dictate synergism.

Recently, many studies have shown that phage-antibiotic combinations are more efficacious than phage-alone treatments [36] [37]. Studies have shown that the synergism between phages and antibiotics varies broadly, and could differ between the antibiotics and phages used [31]. The exact mechanism governing this variation remains unclear. For instance, vancomycin, which works synergistically with one phage, does not act synergistically with other phages or exerts a different intensity of clear with that phage [17], [18]. This indicates that this combination is phage-dependent and unique. Similarly, we found a variation in the treatment effect of the antibiotics used. High synergism was noted with fosfomycin antibiotic, which could be reason out at this point as the effect of inhibition at the initial stage of cell wall synthesis. This is attributed to the effect of the time of action. In other words, inhibition by antibiotics was rapid and started in the very beginning, which enhanced the activity at a very high rate. The oxacillin antibiotic also showed a similar effect as fosfomycin and a higher effect than ciprofloxacin and vancomycin. This adds another important finding that the cell wall synthesis inhibitor was more efficient, except for vancomycin, which remains unanswered at this point. Similarly, our study showed that cell wall synthesis inhibitors are more efficient than DNA synthesis inhibitors. To support our findings, we found much of the study showed that vancomycin does not work in the PAS test. For instance, a study by Dickey et al. showed that vancomycin antibiotics antagonistically act in combination with *S. aureus* phages [13]. Another study by Tkhilaishvili et al. showed no synergism with the vancomycin [38]. However, the efficacy of vancomycin was poor compared to other antibiotics used in this study, and the FIC index in the checkerboard analysis showed that vancomycin was synergistic. Similar to fosfomycin, oxacillin significantly reduces the bacterial burden. We found much of the study that supported our evidence of synergism between oxacillin antibiotic and *S. aureus* phages, a few examples being, a study by Simon et al. showed the high efficiency of oxacillin and phage Sb-1 [39].

In this study, two different mechanisms of action of antibiotics were selected based on the pathogenesis of the organism. Outcome variation among the antibiotic of the same mechanism of action shows that synergisms cannot be easily translated among various antibiotics or the same antibiotic with another phage. Another important factor to be considered in PAS is the treatment order. Three different treatment orders have been widely reported, including simultaneous and staggered by pre- and post-phage treatment [13]. In the current study, we assessed the effect of treatment order in planktonic cultures and compared the efficacy of PRE-, SIM-, and POS-treatment. The treatment orders are known to cause selective pressure on the bacteria in the order of their treatment. Our data demonstrated that the treatment order had a large-fold variation in bacterial reduction. The highest fold reduction was observed for phage, followed by an antibiotic treatment called POS treatment, showing more than a 20-fold increase in planktonic cell reduction when compared to the other two treatment orders. This observation is consistent with the findings of *in vitro* analyses, which showed that staggering treatment (referred to as POS treatment in this study) is more effective than simultaneous treatment [40], [13]. Although *in vitro-based* evidence is frequently reported for these methods, mechanistic interactions that enhance bacterial clearance in the presence of phages and antibiotics and their order of effects are unknown. Many studies have hypothesized that antibiotics can elongate bacterial cells, leading to increased phage production; however, the effect of the treatment order remains unclear [41]. The effective antibiotics in the treatment order were fosfomycin> oxacillin> ciprofloxacin> vancomycin. It was observed that the simultaneous treatment effect had minimal variation among the antibiotics studied. Unlike the large variation observed among the treatment orders within the antibiotics for fosfomycin, ciprofloxacin, and oxacillin, vancomycin did not vary significantly within the treatment order. The change within is as small as 2.42-fold. However, the fold reduction of all treatment orders was higher than the individual treatment effects of phages and antibiotics. Throughout the study, we found that all antibiotics followed a pattern of increased killing effect with phages followed by an antibiotic treatment called POS treatment. At this point, the exact reason for such variation is unexplainable from this study outcome; however, it shades the importance of exploring the molecular aspect of the study to determine the PAS clinical success.

Assessment of the mode of action of the antibiotic with the treatment order again showed that fosfomycin and oxacillin produced the highest effect, followed by ciprofloxacin and vancomycin in all three treatment orders. Interestingly, although vancomycin also targeted the cell wall, it did not produce bacterial eradication similar to the other two cell wall synthesis inhibitors. In conclusion, our study shows that except vancomycin other cell wall synthesis-inhibiting antibiotics had a higher effect than DNA synthesis inhibitors. In general, this could be very specific to the *S. aureus* phage. Furthermore, our study also concludes and leaves a trace to clinical transition that the treatment order plays a major role in the phage-antibiotic synergy treatment outcome.

*In vitro* phage-antibiotic synergy is an important analysis prior to the introduction of treatments into patients. This study reported two primary conclusions: (1) different antibiotics can have varied outcomes even when synergistic, and (2) the order of treatment in PAS is significant. The selection of the most synergistic antibiotics *in vitro* may be a helpful guide during patient treatments, as few combinations can be antagonistic [28]. The phage progeny produced during phage and antibiotics combinatorial treatment should be studied to predict the potential outcomes of the combination during treatment, thus, making such *in vitro* analyses essential before being applied *in vivo*.

## Author Contributions

Conceptualization, A.L., P.M. and R.N.; Methodology, A.L., P.M., and R.N.; Software, A.L.; Validation, A.L., and R.N.; Formal Analysis, A.L. and P.M.; Investigation, R.N.; Resources, R.N.; Data Curation, A.L.; Writing – Original Draft Preparation, A.L.; Writing – Review & Editing, P.M., and R.N.; Funding Acquisition, R.N.” All authors have read and agreed to the published version of the manuscript.

## Acknowledgments

The authors, gratefully acknowledges the Indian Council of Medical Research (ICMR), India, for providing financial assistance (Senior Research Fellowship) to support this research. The authors would like to take this opportunity to thank the management of Vellore Institute of Technology for providing the necessary facilities and encouragement to carry out this work.

## Institutional Review Board Statement

Not applicable

## Informed Consent Statement

Not applicable.

## Data Availability Statement

All data are represented as figures, tables. Any further inquiries may be referred to the corresponding author.

